# Hybridogenetic reproduction of *Pelophylax* water frogs from different R-E hemiclonal population systems from Eastern Ukraine: selective mortality, clonal and ploidy diversity

**DOI:** 10.1101/2024.12.03.626604

**Authors:** Anna Fedorova, Eleonora Pustovalova, Mykola Drohvalenko, Olha Biriuk, Marie Doležálková-Kaštánková, Maryna Kravchenko, Olexii Korshunov, Peter Mikulíček, Lukáš Choleva, Dmytro Holovnia, Dmitrij Dedukh, Dmytro Shabanov

## Abstract

European water frogs from the *Pelophylax esculentus* complex include two sexual species, *P. ridibundus* and *P. lessonae,* and their hybrids, which usually clonally transmit one of the parental species’ genomes. This unique reproductive strategy allows hybrids to reproduce with one or both parental species, creating diverse population systems. Unlike most well-studied population systems in Europe, the Siverskyi Donets River basin in Eastern Ukraine features diploid and polyploid hybrids coexisting with *P. ridibundus*, while *P. lessonae* is absent (R-E systems). To reveal diverse system compositions, genetic divergence, and tadpole selective mortality, we combined novel data from over a decade of observations with previous research on population systems in the Siverskyi Donets River. Two main types of R-E systems were identified: those with diploid hybrids in northern localities and those with both diploid and triploid hybrids, extending from the mainstream of the Siverskyi Donets River to its tributaries. Additionally, we found higher genetic diversity in R-genomes compared to L-genomes, likely due to the absence of *P. lessonae* and the ongoing input of recombined R-genomes from *P. ridibundus* and triploid hybrids. This study highlights the importance of continuous monitoring and research to unravel the dynamics and complexity of water frog population systems.

## Introduction

Hybridization plays a crucial role in the evolution of species, contributing to the creation of new genetic lineages and affecting the genetic diversity of populations (Coyne & Orr, 2009; Schön et al., 2009). However, progenies resulting from interspecies crosses with certain genetic and chromosomal divergence are often sterile (Abbott et al., 2013; Mallet, 2007; Marta et al., 2023). Hybrid sterility can be overcome by several mechanisms. One such mechanism is the transition of hybrids to clonal reproduction, as demonstrated in parthenogenetic and gynogenetic organisms (Dawley & Bogart, 1989; Neaves & Baumann, 2011; Schön et al., 2009; Stöck et al., 2021). Another mechanism involves polyploidization of hybrids, which occurs less often in animals compared to plants (Alix et al., 2017; Schmid et al., 2015; Stöck et al., 2001; Zhou & Gui, 2017) and can be achieved via spontaneous genome duplication or by exploiting clonal reproduction as a transition stage (Stöck et al., 2021). In the latter case, triploids often appear and may serve as a “bridge” towards polyploidization (Husband, 2000). The last mechanism is the transition to hemiclonal reproduction including hybridogenesis. During hybridogenesis, one of the parental genomes is eliminated in germ lines of hybrids, while the other one is endoreplicated leading to the production of haploid clonal gametes (Dawley & Bogart, 1989; Lavanchy & Schwander, 2019; Schön et al., 2009; Stöck et al., 2021). The co-existence of clonal or hemiclonal and polyploid organisms represents a particular interest in their evolution.

Hemiclonal reproduction by hybridogenesis accompanied by polyploidization was found in European water frogs from the *Pelophylax esculentus* complex. This complex comprises parental species *P. ridibundus* (Pallas, 1771), genome RR, and *P. lessonae* (Camerano, 1882), genome LL, together with both diploid (LR) and triploid (LLR, LRR) hybrids *P. esculentus* (Linnaeus, 1758), (Graf & Polls-Pelaz, 1989; Plötner, 2005). Rare tetraploid hybrids (LLRR) have been documented in Swedish and Ukrainian populations (Borkin et al., 2004; Jakob, 2007), and occasional pentaploid hybrid (LLLRR) was obtained in laboratory crosses between frogs from Polish populations (Hermaniuk et al., 2013). To reproduce via hybridogenesis, diploid hybrid frogs usually coexist with one or both parental species (Graf & Polls-Pelaz, 1989; Plötner, 2005) resulting in a variety of population systems with different compositions of parental species and different forms of hybrids (Dufresnes & Mazepa, 2020; Graf & Polls-Pelaz, 1989; Plötner, 2005). Population systems where diploid and sometimes triploid hybrids coexist together with *P. lessonae* individuals (so-called L-E systems) are the most widespread (Hermaniuk et al., 2020; Hoffmann et al., 2015; Hoffmann & Reyer, 2013; Pruvost et al., 2013, 2015). In such systems, hybrids eliminate *P. lessonae* genome from their germline cells, endoreplicate *P. ridibundus* genome, and transmit it to gametes (Dedukh et al., 2019; Tunner & Heppich, 1981; Tunner & Heppich-Tunner, 1991). Gametogenic aberrancies have been rarely observed among males and females from L-E systems (Chmielewska & Kazmierczak et al., 2022; Dedukh et al., 2019). Contrary to L-E, the population systems comprising mostly diploid hybrids and *P. ridibundus* (R-E) are spread in Central and Eastern Europe (Borkin et al., 2004; Doležálková-Kaštánková et al., 2018; Suriadna et al., 2020; Uzzell et al., 1975). Despite the suggestion that hybrids in R-E systems should eliminate *P. ridibundus* genome from their germ line (Graf & Polls-Pelaz, 1989; Uzzell et al., 1977) some hybrids also eliminate *P. lessonae* genome and produce gametes with *P. ridibundus* genome (Pustovalova, et al., 2022a; Ragghianti et al., 2007; Uzzell et al., 1977; Vinogradov et al., 1991). In North Western parts of Europe, “pure” hybrid systems (E systems) were found. In such systems, diploid hybrids produce diploid and haploid gametes as well as triploid hybrids usually form haploid recombined gametes (Christiansen, 2009; Christiansen et al., 2010; Christiansen & Reyer, 2009, 2011; Dedukh et al., 2022; Reyer et al., 2015).

Interestingly, R-E systems strikingly differ in the presence and ratio of frog genotypes and sexes. As Central European ones included only diploid hybrid males coexisting with *P. ridibundus* (Doležálková-Kaštánková et al., 2018), the R-E systems found in Siverskyi Donets River basin (Eastern Ukraine) represent a more complex structure. In the Siverskyi Donets River basin, diploid and polyploid hybrids of both sexes *P. esculentus* coexist with *P. ridibundus.* Moreover, the ratio of parental species and different forms of hybrids (diploid LR, and triploid LLR and LRR) varies across different population systems in this basin (Biriuk et al., 2016; Borkin et al., 2004; Drohvalenko et al., 2022, 2023; Meleshko et al., 2014). Therefore, this region was designated as the Siverkyi Donets center of water frog diversity (Biriuk et al., 2016; D. Shabanov et al., 2020). Despite the long-term study of R-E systems structure, the investigation of R-E systems within the Siverkyi Donets Center of water frogs diversity using a combination of molecular and cytogenetic methods has not yet been undertaken. Thus firstly, by application of these methods, we aim to characterize the population structure of the novel and previously described population systems as well as revise previously published data.

In population systems, where one parental species is absent, its genome is clonally transmitted exclusively by hybrids suggesting its lower genetic diversity (Doležálková-Kaštánková et al., 2018; Pruvost et al., 2015). Earlier, in diploid hybrid males from the Central European R-E systems, the genome of *P. lessonae* was found to be presented as a single hemiclone, which indicated their common origin (Doležálková-Kaštánková et al., 2018). Contrarily, in L-E systems from Central Europe, several clonal lineages of R-genomes were found in hybrids, and all the L- genomes showed variation since they come from *P. lessonae* (Pruvost et al., 2015). In L-E-R and E systems from the same area, clonal lineages of both R- and L-genomes were present indicating the potential multiple emergence of hybrids (Pruvost et al., 2015). Thus, secondly, we aim to determine clonal genomes diversity, along with understanding the mechanisms supporting hybrid reproduction and their origin within systems in the Siverskyi Donets center of water frogs diversity.

All population systems of water frogs are characterized by selective mortality of progeny (Abt Tietje & Reyer, 2004; Binkert et al., 1982). It is supposedly driven by clonal genomes evolution and intergenomic interactions (Christiansen, 2005; Dowling & Secor, 1997; Joly, 2001). Parental species individuals may appear from crosses between hybrids transmitting similar clonal genomes or between hybrids and parental species (Günther & Plötner, 1988). Such individuals, known as hybridolytic, often show developmental abnormalities and usually die before reaching sexual maturity (e.g. Berger, 1968, 1971; Guex et al., 2002; Reyer et al., 2015). Their mortality stabilises water frog reproduction in the population system while maintaining the highest possible hybrid ratio (Reyer et al., 2015; Vorburger, 2001). Individuals of parental species were frequently found among offspring as hybrid males and females from the Siverkyi Donets River basin produce gametes with R- and L- genomes exclusively or simultaneously (a phenomenon of “hybrid amphigameticity”) (Biriuk et al., 2016; Dedukh et al., 2017, 2022; A. Fedorova & Shabanov, 2022). Moreover, they produce a low number of gametes with LL, RR, and LR genomes as well as a portion of aneuploid gametes (Biriuk et al., 2016; Dedukh et al., 2015, 2017; Pustovalova, et al., 2022a; Pustovalova et al., 2024). Such a variety of gametes produced by hybrids leads to the divergence of genome composition in the progeny while it is unclear which genotypes can survive. Therefore, our third aim was to analyze the selective mortality in selected population systems within the Siverskyi Donets center of water frogs diversity.

In a current study, we provide summarized results obtained during more than a decade of observations by different methods on population systems of water frogs from the Siverskyi Donets River basin by analyzing: 1) genotypic compositions of water frogs in various population systems; 2) genetic diversity and clonal lineages of R- and L-genomes; 3) selective mortality of the offspring in selected population systems.

## Materials and Methods

### Sampling and data collection

From 2014 to 2024, we caught and analyzed 1888 adult frogs from 18 waterbodies in Eastern Ukraine (Fig. 1; Table 1; Table S2). Fourteen waterbodies belonging to the Siverskyi Donets River basin were the main area of study (in total, 1848 adult frogs); and 40 frogs from the remaining four waterbodies belonging to the Dnipro River basin were used as an outgroup for population-genetic analysis (Table 2; Table S2). The localities studied within the Dnipro River basin belonged to basins of four tributaries of Dnipro (Seym River, Udai River, Psel River, and Merla River), but for simplicity, all are referred to as “Dnipro basin” through the paper. The names, abbreviations, and geographical coordinates of the studied localities are presented in Table 1.

**Figure 1.**
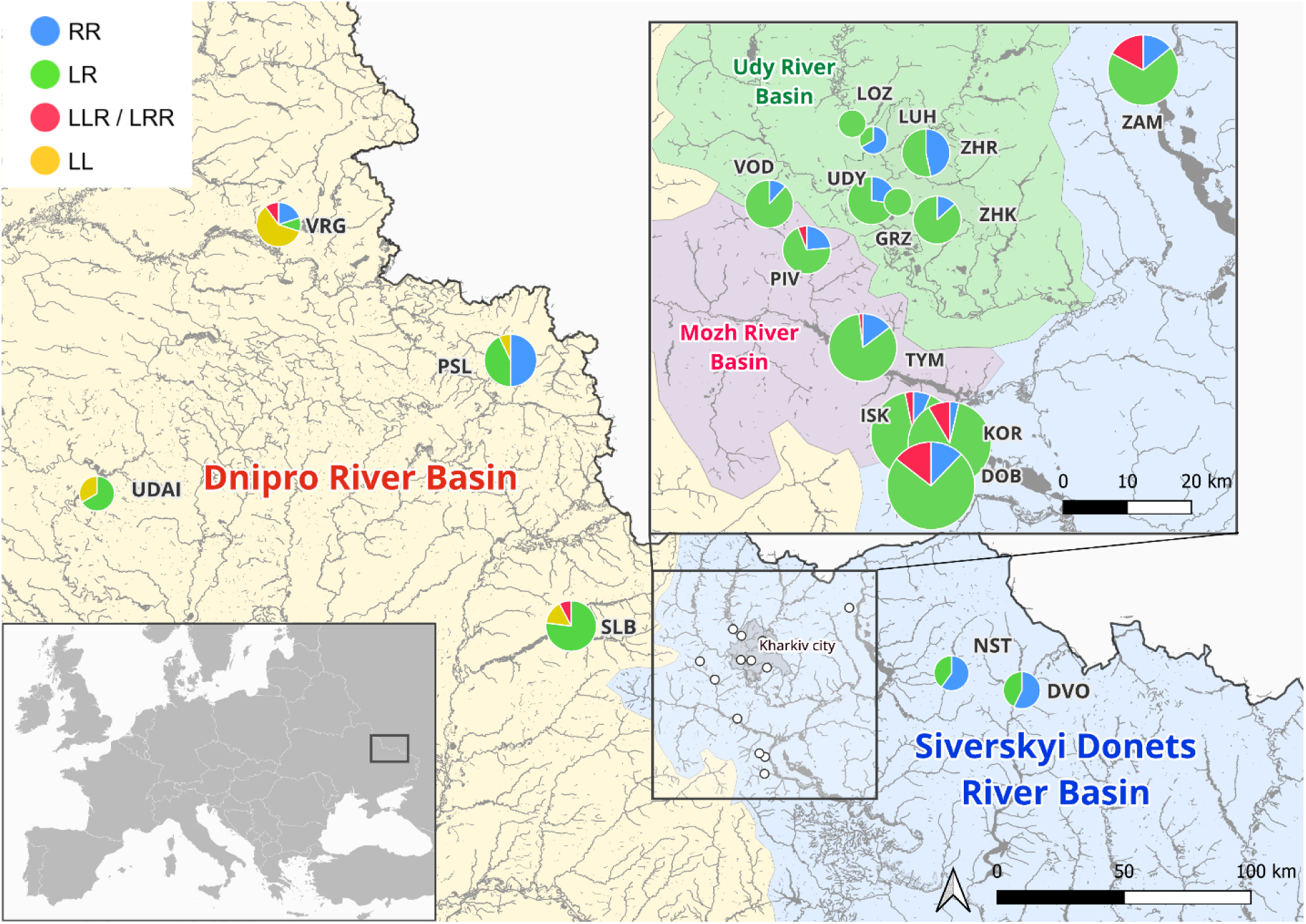
Sampling sites of water frogs and compositions of HPSs. DOB – Lower Dobrytskyi pond, DVO – NNP “Dvorichanskyi”, GRZ – Gruzotonovka, ISK – Iskiv pond, KOR – Koriakiv pond, LOZ – Lozovenky, LUH – Oleksiivskyi Luhopark, NST – Nesterivka, PIV – Pivdenne, PSL – Psel river, SLB – NNP “Slobozhanskyi”, TYM – Tymchenky pond, UDAI – Udai river, UDY – Udy river, VOD – Vodiane, ZAM – Zamulivka village, ZHK – Zhykhorets river, ZHR – Zhuravlivskyi Hydropark. The size of the pie-charts corresponds with the number of frogs analyzed ranging from two frogs in GRZ to 770 frogs in DOB (see details in Table 1).

**Table 1.**
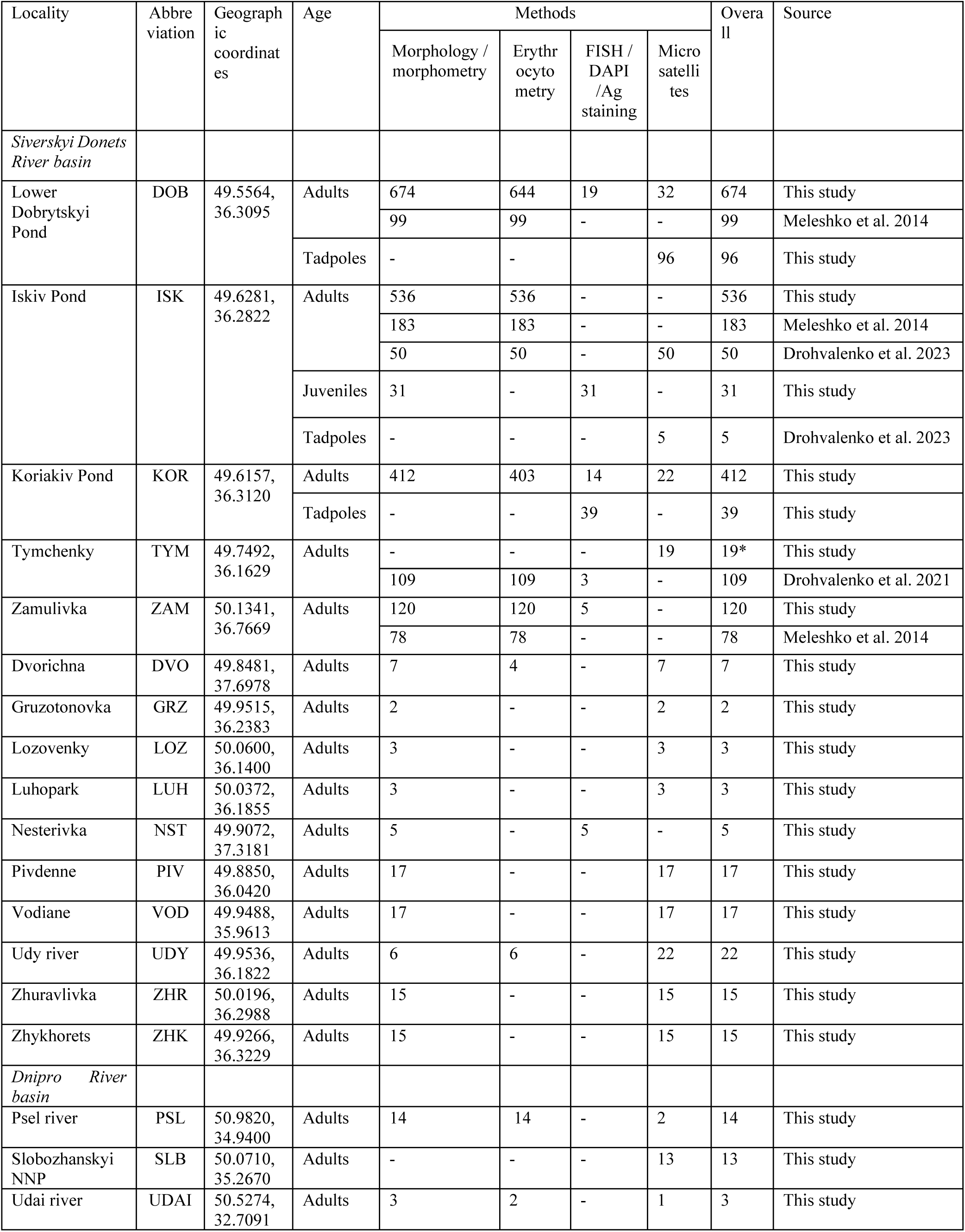
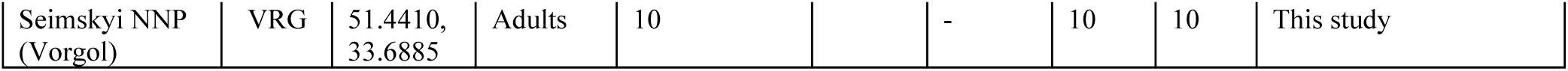
Methods and numbers of frogs used for the description of 18 HPSs in Siverkyi Donets River and Dnipro River basins. * - same frogs as in Drohvalenko et al. 2021

**Table 2.** Distribution of *Pelophylax* species in studied localities and types of HPSs based on compositions of adult frogs. Combined data from Meleshko et al. 2014, Drohvalenko et al. 2021, and this study.

| Locality | HPS type | RR | LR | LL | LRR | LLR | LLR/LRR | Overall | Source |
| --- | --- | --- | --- | --- | --- | --- | --- | --- | --- |
| *Siverskyi Donets River basin* |  |  |  |  |  |  |  |  |  |
| DOB | R-E-Ep | 90 | 563 | - | 11 | 2 | 107 | 773 | This study<br>Meleshko et al. 2014 |
| ISK | R-E-Ep | 48 | 696 | - | 6 | 2 | 17 | 769 | This study<br>Meleshko et al. 2014<br>Drohvalenko et al. 2023 |
| KOR | R-E-Ep | 15 | 365 | - | 2 | 4 | 26 | 412 | This study |
| TYM | R-E-Ep | 16 | 91 | - | - | 2 | - | 109 | This study<br>Drohvalenko et al. 2021 |
| ZAM | R-E-Ep | 28 | 136 | - | - | - | 34 | 198 | This study<br>Meleshko et al. 2014 |
| DVO | R-E | 4 | 3 | - | - | - | - | 7 | This study |
| GRZ | R-E | - | 2 | - | - | - | - | 2 | This study |
| LOZ | R-E | - | 3 | - | - | - | - | 3 | This study |
| LUH | R-E | 2 | 1 | - | - | - | - | 3 | This study |
| NST | R-E | 3 | 2 | - | - | - | - | 5 | This study |
| PIV | R-E-Ep | 4 | 12 | - | 1 | - | - | 17 | This study |
| VOD | R-E | 2 | 15 | - | - | - | - | 17 | This study |
| UDY | R-E | 6 | 16 | - | - | - | - | 22 | This study |
| ZHR | R-E | 7 | 8 | - | - | - | - | 15 | This study |
| ZHK | R-E | 2 | 13 | - | - | - | - | 15 | This study |
| *Dnipro River basin* |  |  |  |  |  |  |  |  |  |
| SLB | L-E-Ep | - | 10 | 2 | 1 | - | - | 13 | This study |
| VRG | L-E-Ep-R | 2 | 1 | 6 | 1 | - | - | 10 | This study |
| PSL | L-E-R | 7 | 6 | 1 | - | - | - | 14 | This study |
| UDAI | L-E | - | 2 | 1 | - | - | - | 3 | This study |

Sampling was carried out at night by hand using a flashlight or during the day with a dip net. Additionally, 140 tadpoles and 31 froglets (juvenile individuals after metamorphosis but before the first hibernation) were collected from three localities in the Siverskyi Donets River basin (Table 1).

For further analysis of population systems composition, we combined our data on 1888 frogs from 14 population systems with previously published data on 519 frogs from four out of 14 population systems in the Siverskyi Donets River basin (Drohvalenko et al., 2022, 2023; Meleshko et al., 2014) (Table 1, 2).

To emphasize the complex composition of frogs’ genotypes in different localities within the Siverskyi Donets River basin, we use the term “hemiclonal population system” (HPS) and signify the presence of *P. ridibundus* by R, diploid hybrids by E and polyploid hybrids by Ep (Günther, 1975; D. Shabanov et al., 2020). For example, “R-E-Ep HPS” would stand for hemiclonal population system consisting of *P. ridibundus*, and diploid and triploid *P. esculentus*. Here and throughout the paper, we will use “HPS”, “system”, and “population system” as synonyms. Maps were created using QGIS 3.34.1-Prizren.

Animal collection and further manipulations were approved by The Committee on Bioethics of Karazin National University, Kharkiv, Ukraine (No. 4/16, April 21, 2016; No. 1/23, March 15, 2023). Additionally, we adhered to the Guidelines for Use of Live Amphibians and Reptiles in Field and Laboratory Research, the Ukrainian Law on the Protection of Animals from Cruelty, and international ARRIVE guidelines.

### Taxonomical identification by morphology

Initially, to differentiate between hybrid *P. esculentus* and its parental species *P. ridibundus* and *P. lessonae,* we thoroughly examined four main morphological traits of each frog: the shape of the metatarsal tubercle, the coloration of the inguinal region underside of thighs, the coloration of vocal sacs, and the spotting pattern (Table S1). A combination of these traits allows reliable taxonomical identification of water frogs from the *Pelophylax esculentus* complex (Berger et al., 1978; Günther, 1975; Kierzkowski et al., 2011). For tadpoles, we identified developmental stages following Gosner, (1960).

### Ploidy and genotypic identification by erythrocyte cytometry, Ag-, DAPI-stainings and FISH

To distinguish between diploid and triploid hybrids we took air-dried blood smears from most of the adult individuals (n=1713) and measured erythrocyte lengths (Table S2) following (Ogielska-Nowak, (1978) and Bondarieva et al., (2012). Erythrocytes were photographed and measured under ×40 magnification with either Leica DM 2000 microscope or a Carl Zeiss Jena Amplival microscope.

To obtain chromosomal preparations, we followed the protocol described by Pustovalova et al. (2022a). We injected adults (n=46) and froglets (n=31) with 0.04% colchicine solution peritoneally and leaving them for 12-24 hours; tadpoles (n=131) were placed in a 0.4% colchicine solution for 12 hours, before being sacrificed. Dissected tissues (bone marrow, gonads, and intestines from adults and froglets, and gills, regenerated tail and intestines from tadpoles) were hypotonized in 0.07M KCl for 20-30 minutes, fixed in Carnoy’s solution (3 methanol: 1 acetic acid) and stored at +4 °C. Sex identification of tadpoles was based on the analysis of gonadal morphology (Christiansen, 2009; Haczkiewicz & Ogielska, 2013).

To identify the ploidy of 264 individuals more precisely (88 froglets, 140 tadpoles, 36 adults) we applied Ag^+^-staining to reveal nucleolus organizer regions (NORs) in nuclei and chromosomes following the protocol described by (Howell & Black, 1979). Individuals with 26 chromosomes (2n=26) and two nucleoli in nuclei were identified as diploid (Fig. S1, A, B); individuals with 39 chromosomes (3n=39) and three nucleoli in nuclei were identified as triploid (Fig. S1, C, D).

To clearly identify genome composition, we performed fluorescent *in situ* hybridization (FISH) with two species-specific probes on mitotic chromosomes from somatic tissues of 39 tadpoles and 31 froglets according to protocol described by (Choleva et al., 2023). For *RrS1* (labeled with biotin (Roche)) we used the following primers (Ragghianti et al., 2007):

Forward: 5’-AAGCCGATTTTAGACAAGATTGC-3’
Reverse: 5’-GGCCTTTGGTTACCAAATGC-3’

For *PlesSat01-48* (labeled with digoxigenin (Roche)) we used the following primers (Choleva et al., 2023):

Forward: 5’-TTTGGCTTCCAAGGGCCGGG-3
Reverse: 5’-TGACCAAAAACGACACTCCC-3

The mixture contained 3 ng/μL of *RrS1* and *PlesSat01-48* probes was denatured at +86 °C for 10 minutes and applied on chromosomes previously denatured at +75 °C in 75% formamide and 2**×** SSC. After 12-15 hours of hybridization, probes were detected using streptavidin-Alexa488 (S11223, Invitrogen) and anti-digoxigenin-rhodopsin (11207750910, Merck) for 3-4 hours. Slides were mounted in fluoroshield with DAPI. Additionally, we used DAPI-staining of pericentromeric loci to identify the genome composition of tadpoles (n=39) following Pustovalova et al. (2022b). Slides were photographed using an Olympus Provis BX53 microscope with a CCD camera (DP30W Olympus) and a Leica DM2000 microscope with a DFC3000 G camera, both equipped with standard fluorescent filter sets. For each individual, we photographed all countable mitotic chromosomes and manually counted the number of chromosomes and pericentromeric loci after DAPI-staining. The pericentromeric loci on *P. ridibundus* chromosomes show a stronger signal due to the higher number of tandem repeats compared to *P. lessonae* (Marracci et al., 2011; Ragghianti et al., 1995), allowing us to distinguish between the two species. Chromosomes with strong signals at all loci were identified as *P. ridibundus*, while those with half showing weaker signals were marked as *P. lessonae*. For triploid hybrids, genomes were classified as LRR if one-third of the chromosomes lacked strong signals and LLR if two-thirds did (Fig. S2, A, C, E, G). For FISH analysis, we manually counted the number of *RrS1* and/or *PlesSat01-48* signals in each mitosis. Mitosis with only *RrS1* signals indicated *P. ridibundus* chromosomes. Hybrids and their ploidy were identified by the presence of two (LR, LRR) or four (LLR) *PlesSat01-48* signals, and 13 (LR, LLR) or 26 (RRL) *RrS1* signals (Fig. S2, B, D, F, H).

### Population genetic analysis based on microsatellite loci

A part of the analyzed individuals (193 adults, 96 tadpoles) was also used for microsatellite analysis for precise identification of genotypes, further population genetic analysis, and identification of clonal lineages.

Genomic DNA was extracted from blood, fingertips (for adults), or tale tips (for tadpoles) stored in 96% ethanol using the NucleoSpin Tissue kit (Macherey-Nagel, Düren, Germany) or Geneaid DNA Isolation Kit (Geneaid, New Taipei City, Taiwan) following manufacturers protocols.

A total of 193 adult frogs were genotyped with 14 microsatellite loci amplifying in L- and/or R- genomes. Microsatellite markers were amplified using Multiplex Hot-Start PCR Master Mix, 2x (Biotechrabbit, Berlin, Germany). All the loci were grouped in two multiplex PCR sets.

Multiplex 1: Re1Caga10 (Arioli et al., 2010), Ga1a23, Rrid013A (Christiansen & Reyer, 2009), Rrid171A (Hotz et al., 2001), RICA18, RICA1b5, RICA1b6 (Garner et al., 2000).

Multiplex 2: Res17 (Zeisset et al., 2000), GA1A19 (Arioli et al., 2010; Pruvost et al., 2013), RICA2a34 (Christiansen & Reyer, 2009), RICA5, RICA1b20 (Garner et al., 2000), Rrid082A (Hotz et al., 2001), Re2Caga3 (Arioli et al., 2010).

Additionally, 96 tadpoles were genotyped with 18 microsatellite loci amplified in L- and/or R- genomes. Microsatellite markers amplification for tadpoles was carried out with a 2x Qiagen multiplex PCR kit (Qiagen, Valencia, CA). All the loci were grouped in four multiplex PCR sets. Multiplex 1: GA1A19 (Arioli et al., 2010; Pruvost et al., 2013), RlCA18, RlCA1b5 (Garner et al., 2000), RlCA2a34 (Christiansen & Reyer, 2009), Rrid013A (Hotz et al., 2001).

Multiplex 2: Re1Caga10, Re2Caga3 (Arioli et al., 2010), Rrid059A (Christiansen & Reyer, 2009; Hotz et al., 2001).

Multiplex 3: RlCA5 (Garner et al., 2000), Rrid082A (Hotz et al., 2001), Res17, Res22 (Zeisset et al., 2000).

Multiplex 4: Rrid171A (Hotz et al., 2001), CA1a27, Ga1a23, Rrid064A, Rrid135A (Christiansen & Reyer, 2009).

Alleles were scored with GeneMarker® software (SoftGenetics, USA).

To evaluate genetic diversity we estimated the mean number of alleles per locus (Na) and expected heterozygosity (He). To estimate the clonality of R- and L-genomes we analyzed multilocus genotypes (MLGs), which are defined as the identical sets of alleles in the analyzed microsatellite loci. The presence of identical MLGs is considered an indication of clonal reproduction. However, the same MLGs could also be found in individuals with sexual reproduction if selected molecular markers have low discrimination power. To estimate the discrimination power of selected set of microsatellite loci, we calculated the probability of identity (PI) (Pruvost et al., 2015). The estimated PIs for eight *P. lessonae*, and 40 *P. ridibundus*, were 1.3×10^-5^, and 7.8×10^-7^, respectively. Both PIs are very low, showing that the selected set of microsatellite loci is informative for MLG analysis and estimation of clonality.

All the population-genetic indexes and MLGs were calculated with GenAlEx 6.5 package for MS Excel (Peakall & Smouse, 2012).

## Results

### Compositions and dynamics of HPSs

The analysis of new (n=1848) and earlier published animals (n=519, (Drohvalenko et al., 2022, 2023; Meleshko et al., 2014) from 14 HPS in the Siverskyi Donets River basin revealed 227 *P. ridibundus* and 2140 *P. esculentus* (1926 diploids and 214 triploids), with no *P. lessonae* registered (Table 2).

In the Siverskyi Donets basin, we identified eight R-E HPS and six R-E-Ep HPS (Fig. 1, Table S2). Diploid *P. esculentus* prevailed in 13 out of 14 HPSs (Fig. 1; Table 2; Table S2). In all but two (LOZ, GRZ) localities we found *P. ridibundus*. However, we suggest that these two HPSs are also inhabited by *P. ridibundus* as our samples were insufficient (n=2, n=3 accordingly). Considering the presence of *P. ridibundus* in all the surrounding HPS (Fig. 1), we assume that these HPSs belong to the R-E-type. Triploids were found in the main streambed of the Siverskyi Donets River and its tributary (the Mozh River) (ranging from 2% to 14%). Among triploids, both LRR and LLR genotypes were present (Table 2). In other tributaries (the Udy River and the Oskil River), we found only diploid *P. esculentus* (Fig. 1).

Among 40 frogs from the Dnipro River basin, nine were *P. ridibundus*, 21 were *P. esculentus* (19 diploids and 2 triploids), and 10 were *P. lessonae* (Table 2).

The occurrence and proportions of different forms of frogs changed over time in five R-E-Ep HPSs (Fig. 2, Table S2). Diploid *P. esculentu*s was the prevailing form in all studied HPS each year, ranging from 51% in ZAM to 100% in ISK (Fig. 2). HPS in ZAM had the highest proportion of triploids, up to 29% in 2013 (Fig X). The proportion of *P. ridibundus* in all five HPS ranged from 0% to 23% (Fig. 2). The proportions of different forms of frogs changed significantly through the years (Fig. 2).

**Figure 2.**
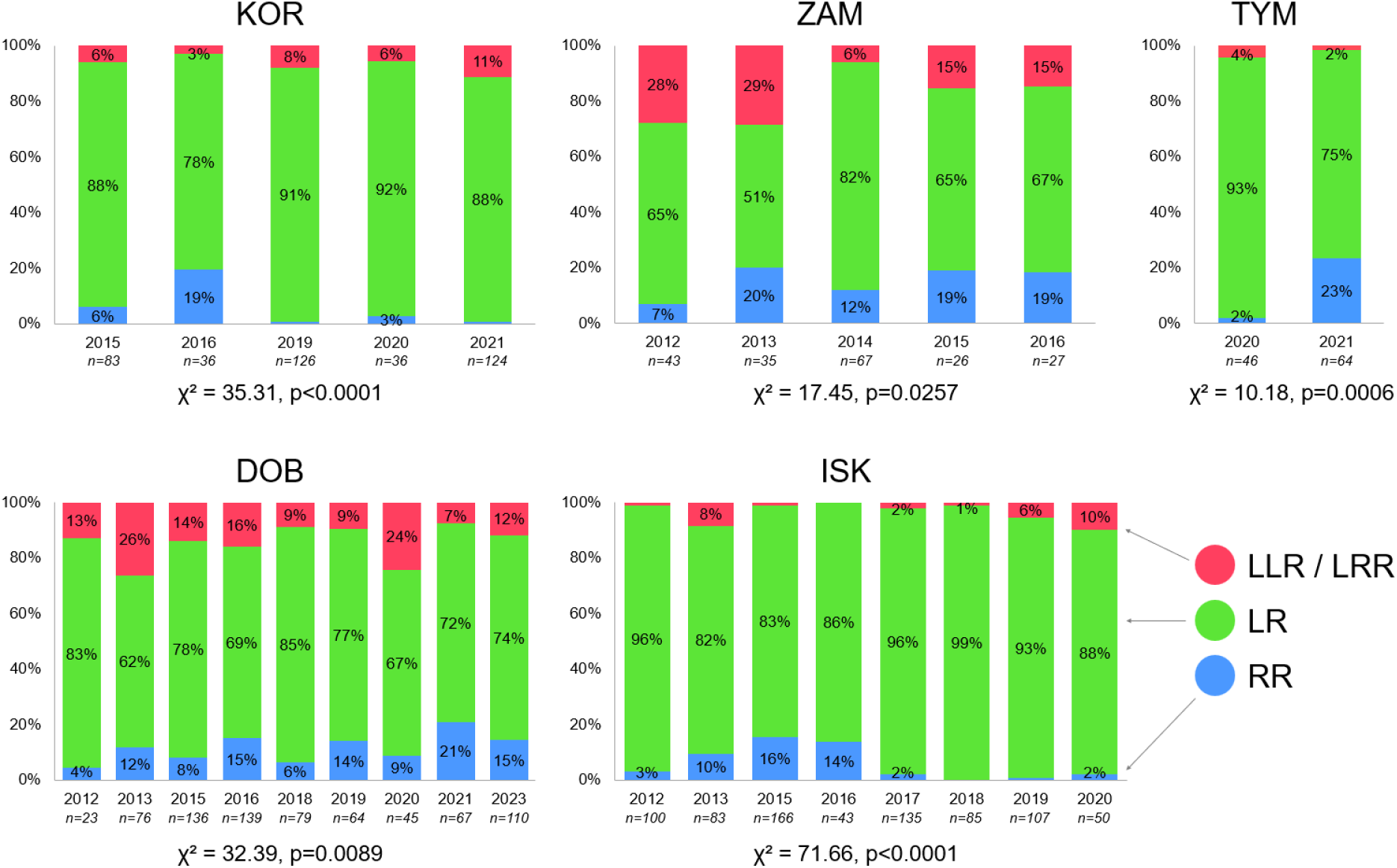
Composition and dynamics of five HPS. Combined data from Meleshko et al. 2014, Drohvalenko et al. 2021, and this study. DOB - Lower Dobrytskyi pond, ISK - Iskiv pond, ZAM - Zamulivka, KOR - Koriakiv pond, TYM - Tymchenky.

Our study examines the temporal changes in the occurrence and distribution of various frog forms across five R-E-Ep HPSs (Fig. 2, Table S2). Over nine and eight years of monitoring HPS composition in DOB and ISK, respectively, we consistently found diploid *P. esculentus* to be the dominant form, ranging from 62% to 85% in DOB and from 82% to 96% in ISK annually. Triploid individuals ranged from 7% to 26% in DOB and from 0% to 10% in ISK. Additionally, the proportion of *P. ridibundus* varied from 4% to 21% in DOB and from 0% to 16% in ISK.

In a five-year study of KOR and ZAM, diploid *P. esculentus* similarly prevailed, accounting for 78% to 92% in KOR and 51% to 82% in ZAM. Triploid *P. esculentus* ranged from 3% to 11% in KOR and from 6% to 29% in ZAM. *P. ridibundus* varied from 0% to 19% in KOR and from 7% to 20% in ZAM. Detailed data on the population system composition of DOB, ISK, KOR, and ZAM for each year can be found in Fig. 2. Observations spanning two years in TYM were documented elsewhere (Drohvalenko et al., 2022). As for the other nine localities, they were each investigated only once, precluding a comprehensive description of the dynamics of HPS composition in those areas.

### Genetic diversity

For 193 frogs from 15 localities (11 from Siverskyi Donets and four from Dnipro, Table 1), genetic diversity and clonal lineages were estimated using microsatellite analysis. From the initial 14 microsatellite loci, we used 12 for further analysis while two were excluded (Re1Caga10 amplified in 11% of samples only; RICA1b20 had alleles that were non-specific, i.e. the same alleles amplified in both L- and R-genomes) (Table S3). All 12 microsatellite loci were polymorphic among all the studied individuals (Table S3). The loci RICA18, Ga1a23, RICA2a34, and RICA5 amplified in L-genomes, while the loci Res17, Rrid082A amplified in R-genomes. Loci Rrid171A and Re2Caga3, which are typically R-specific (Arioli et al., 2010; Hotz et al., 2001), also amplified in 75% and 38% of *P. lessonae* individuals, respectively. The remaining loci (RICA1b5, RICA1b6, Rrid013A, GA1A19) were bi-specific (the loci amplified in both L- and R-genomes but different alleles were found in either genomes).

Triploidy was identified by the presence of three alleles in the same locus or the allele-dosage effects, and additionally supported by erythrocyte cytometry and karyoanalysis.

In total, in L- and R-genomes we found 37 and 77 microsatellite alleles, respectively (Table S3). The estimated population genetic indexes for L- and R-genomes are presented in Table 3. For both L- and R-genomes from the Dnipro basin both Na (average number of alleles per locus) and He (unbiased expected heterozygosity) indexes were almost similar, with L-genomes having both indexes only ∼1.1 times lower (p=0.7914) (Table 3). Alternatively, in the Siverskyi Donets basin, L-genomes had ∼4 times lower Na and ∼2.8 times lower He (p=0.0086) (Table 3) suggesting a much lower diversity of L-genomes in studied HPSs.

**Table 3.** Population genetic indexes for L- and R-genomes from two river basins. Na - the average number of alleles per locus. He - unbiased expected heterozygosity.

|  | N of HPSs | N of samples | Na | He±SE |
| --- | --- | --- | --- | --- |
| *L-genomes* |  |  |  |  |
| Siverskyi Donets basin (all HPS) | 12 | 131 | 2.4 | 0.190±0.080 |
| Siverskyi Donets basin (R-E HPS) | 8 | 61 | 2.25 | 0.222±0.079 |
| Siverskyi Donets basin (R-E-Ep HPS) | 4 | 70 | 1.88 | 0.153±0.080 |
| Dnipro basin | 4 | 22 | 4.4 | 0.416±0.126 |
| All | 16 | 153 | 4.7 | 0.259±0.082 |
| *R-genomes* |  |  |  |  |
| Siverskyi Donets basin | 12 | 168 | 9.5 | 0.542±0.073 |
| Siverskyi Donets basin (R-E HPS) | 8 | 83 | 8.0 | 0.546±0.068 |
| Siverskyi Donets basin (R-E-Ep HPS) | 4 | 85 | 7.1 | 0.518±0.075 |
| Dnipro basin | 3 | 17 | 4.7 | 0.456±0.072 |
| All | 15 | 185 | 10.6 | 0.551±0.072 |

### Clonal lineages

Based on the data provided by microsatellite analysis we identified clonal lineages of both R- and L-genomes by estimating their MLGs.

In all three LLR and two LL (out of eight LL) individuals having only one heterozygous locus, we could distinguish between two different L-genomes. Therefore, we analyzed the respective two L-genomes separately. For the individuals who had two or more heterozygous loci, we were not able to separate two different genomes and, therefore, we analyzed them as one MLG. Among 158 L- genomes in 153 frogs (LL, LR, LLR, and LRR), we have found 52 MLGs. Seventeen MLGs, and therefore clonal lineages, were shared among a minimum of two and a maximum of 27 individuals while the remaining 37 MLGs were unique (Table S4). All the clonal lineages among L-genomes were found in frogs from the Siverskyi Donets basin, while all the MLGs of frogs from the Dnipro basin were unique (Table S4).

Among 10 LRR and 40 RR frogs, we found only two LRR individuals with only one heterozygous locus allowing us to distinguish between two different R-genomes. All the remaining 50 frogs had two or more heterozygous loci. For these frogs, we could not separate two different genomes and therefore analyzed them as one MLG. Among 187 R-genomes in 185 frogs (RR, LR, LLR, and LRR), we found 180 MLGs. One MLG was shared among five individuals, and three were shared among two individuals each. These MLGs, and therefore clonal lineages, were found in frogs from both Siverskyi Donets and Dnipro basins (Table S4). The remaining 176 MLGs were unique (Table S4).

### Genetic composition and selective mortality of tadpoles and froglets from three HPSs

We analyzed the genetic compositions of tadpoles from two localities, DOB (96 individuals) and KOR (39 individuals), as well as froglets from ISK locality (31 individuals). Among all the tadpoles, we found 75 *P. ridibundus*, 44 diploid *P. esculentus*, and 16 triploid *P. esculentus* (13 with LRR genotype and three with LLR genotype). Among 31 froglets, 21 were *P. ridibundus*, and 10 were diploid *P. esculentus.* No *P. lessonae* tadpoles or froglets were detected.

For 96 tadpoles from DOB, genotypes were identified with microsatellite analysis. In total we analyzed 17 polymorphic loci, among which five were L-specific (RlCA18, RICA2a34, RlCA5, CA1a27, Ga1a23), seven were R-specific (Re2Caga3, Rrid082A, Res17, Rrid064A, Rrid135A, Res22) and six amplified in both L- and R-genomes (GA1A19, RlCA1b5, Rrid013A, Re1Caga10, Rrid059A, Rrid171A). We did not observe the allele-dosage effect in triploids. Mean number of alleles was 3.0 and 5.9 for L- and R-genomes, respectively. We found 64 *P. ridibundus* and 32 *P. esculentus* (24 LR, seven LRR, and one LLR, sex was not identified).

For 39 tadpoles from KOR, genotypes were identified with DAPI-staining of chromosomes and further confirmed by FISH with species-specific probes. Using analysis of at least five full mitotic metaphases, we revealed 20 diploid LR hybrids (11 males, three females, and six individuals with unknown sex) (Fig. S2, A, B), 11 *P. ridibundus* tadpoles (two males, six females, and three individuals with unknown sex) (Fig. S2, C, D), eight triploids (two LRR males, five LRR females (Fig. S2, E, F) and one LLR male (Fig. S2, G, H)).

For 31 froglets from ISK, genotypes were identified using karyotyping followed by FISH with species-specific probes. Applying analysis of at least five full mitotic metaphases, we revealed 21 *P. ridibundus* (all females) and 10 diploid *P. esculentus* (eight males and two females) froglets (Table S2). Additionally, five tadpoles from ISK were analyzed with the same set of microsatellite loci as tadpoles from DOB. Among these five tadpoles, one was *P. ridibunuds* and four were diploid *P. esculentus* (Table S2).

The comparison of proportions between frog genotypes among tadpoles or froglets and adults from DOB, KOR, and ISK revealed that the genetic compositions of the young generation significantly differed from adults (Fig. 3). In all studied localities, the proportion of *P. ridibundus* individuals decreased significantly among adult individuals compared to tadpoles and froglets (p<0.0001). In KOR we also registered a significant difference in proportions of triploids (p=0.0172), while in the other two systems, the difference was not significant (ISK: p=0.6216; DOB: p=0.0850).

**Figure 3.**
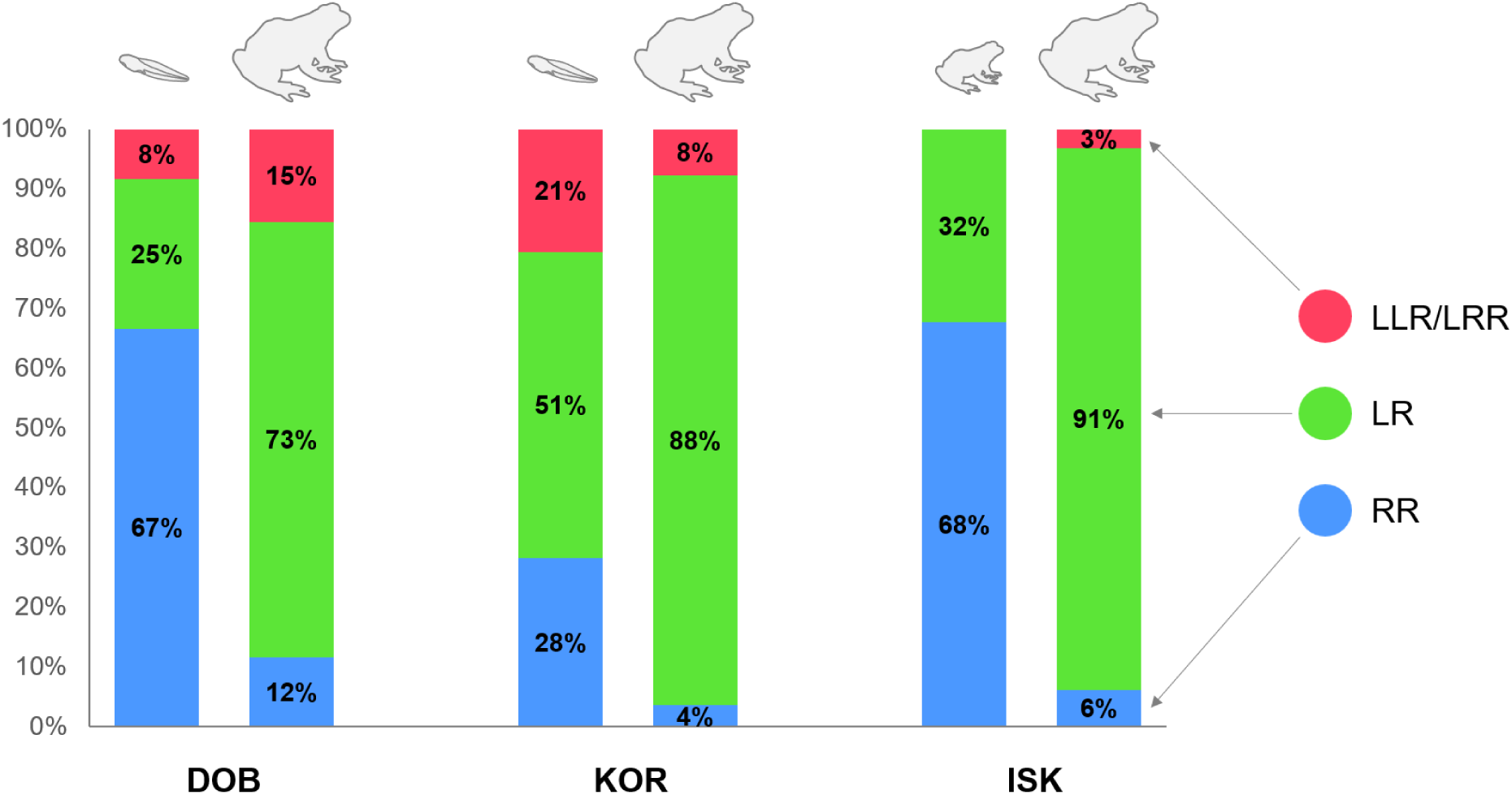
Compositions of adult frogs, froglets, and tadpoles in three HPSs. DOB – Lower Dobrytskyi pond, ISK – Iskiv pond, KOR – Koriakiv pond.

## Discussion

After 12 years of intensive study, we have summarized comprehensive data on the distribution of water frog species, their diversity, and compositions of hemiclonal population systems in the Siverskyi Donets River basin. We investigated water frogs from 10 new localities and increased the number of samples for four localities, which were analyzed before (Drohvalenko et al., 2022, 2023; Meleshko et al., 2014). Moreover, we assessed genetic divergence and clonal lineage distribution in hybrid water frogs along with the selective mortality of tadpoles, which were not studied before in the Siverskyi Donets River basin.

### HPSs compositions and dynamics in the Siverskyi Donets basin

Our results show that in the Siverkyi Donets basin, all the studied HPSs included parental species *P. ridibundus* and hybrid *P. esculentus*. The hybrid taxon was the prevailing form in the majority of HPSs. Moreover, apart from diploid hybrids, we also found a richness of LLR and LRR hybrids in six studied HPSs. Such richness is possible due to both diploid and triploid hybrids producing the diversity of gametes which thus provide the HPSs prosperity. Therefore, we defined two types of R-E systems in the Siverkyi Donets basin, namely, R-E HPS, comprising *P. ridibudnus* and diploid hybrids, and R-E-Ep HPS, comprising *P. ridibundus* and diploid and triploid hybrids. The *P. ridibundus* and triploid hybrids bring valuable contribution into self-sustaining equilibrium of these systems. Earlier, it was considered that R-E-Ep HPSs could be found only in the mainstream of Siverskyi Donets River and nearby water bodies (Borkin et al., 2004; D. Shabanov et al., 2020). The tributaries, such as the Mozh River and the Udy River were suggested to be inhabited only by R-E HPSs (Borkin et al., 2004; D. Shabanov et al., 2020). However, we have shown that triploid hybrids were present in the Mozh River basin (Drohvalenko et al., 2022) and spread up to its tributaries (this study, Fig. 1). In the nearby localities belonging to the Udy River basin, we did not find any triploids, which would suggest the presence of R-E-Ep HPS (Fig. 1).

Based on our analysis of HPSs in the Siverkyi Donets basin and previously obtained data (Borkin et al., 2004; Drohvalenko et al., 2022, 2023; Meleshko et al., 2014; D. Shabanov et al., 2020), we describe three distinct regions of water frog diversity (Fig. 4). The northern R-E region is predominantly located in the Udy River basin (Fig. 4, green). Moving southward, there is the R-E-Ep region along the Siverskyi Donets River and its tributaries hosts triploid hybrids (Fig. 4, red). To the east, the R-Epf region accommodates both male and female *P. ridibundus* alongside exclusively female triploid *P. esculentus* (Borkin et al., 2004; Drohvalenko et al., 2017) (Fig. 4, purple).

**Figure 4.**
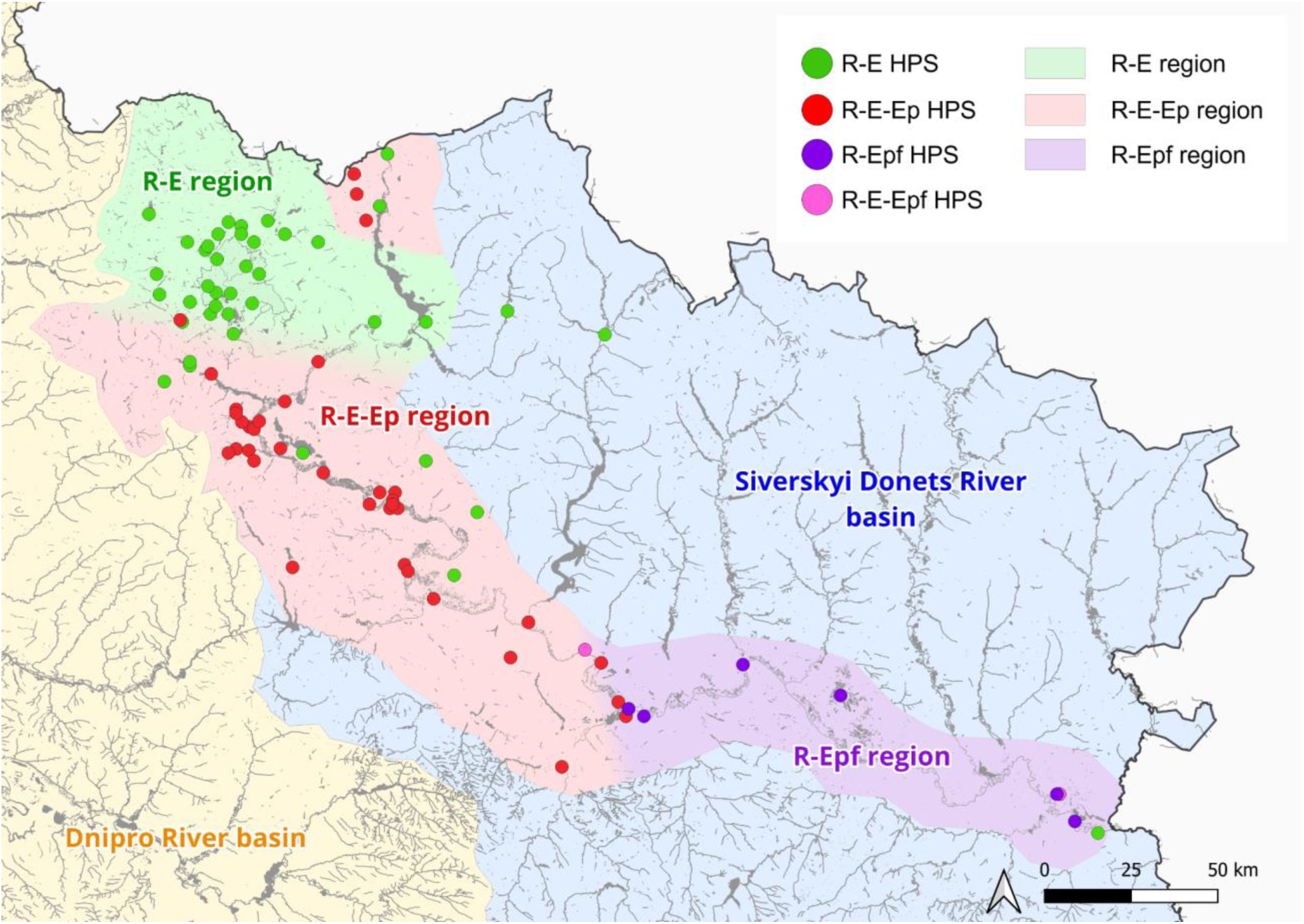
Siverskyi Donets center of diversity of water frogs. Combined data from Biriuk et al., 2016; Borkin et al., 2004; Drohvalenko et al., 2017; D. A. Shabanov, 2014, and this study.

We revealed fluctuations in genetic composition and taxon ratios over time within the studied population systems. Proportions of different forms have been changing significantly through the years (Fig. 2). However, within past years these HPSs tended to remain more stable with diploid hybrid males being the prevailing form. In ISK we even observed a decrease in *P. ridibundus* within the past years. Earlier research has suggested that the anticipated consistency in water frog population systems over time may be attributed to taxon-specific habitat preferences, variations in larval development according to habitat, individual site fidelity, and high annual survival rates (Dufresnes & Mazepa, 2020; Plötner, 2005; Reyer et al., 2015). Another factor playing an important role is the influence of genetic material introgressions from other *Pelophylax* species, which may affect hybridogenetic reproduction by altering gametogenesis in hybrids (Pustovalova et al., 2024). As a result, it may lead to the variability of genome composition and ploidy level of gametes produced by hybrids, and thus change the proportion of genotypes in investigated HPSs (Pustovalova et al., 2024). Significant increase in proportions of *P. ridibundus* and *P. esculentus* with a simultaneous decline in *P. lessonae* was shown in several HPSs from Poland over the last 50 years (Kolenda et al., 2024). Some of these systems completely changed their composition, from L-E or L-E-R to R-E. Moreover, in some water frog population systems in Poland, the introgressions of *P. kurtmuelleri* genome were found (Kolenda et al., 2017).

Assortative mating, female fecundity, and competition among tadpoles and juveniles are crucial for maintaining the stable coexistence of different water frog forms (D. Shabanov et al., 2020; Som et al., 2000). Conversely, fluctuations in water levels, associated with habitat changes, impact the structure and dynamics of water frog populations near rivers (Plötner, 2001). Determining changes in genetic composition and taxon ratios in HPSs is challenging due to varying habitat conditions, limited sample sizes, and sampling errors. Stochastic processes like water drying and multiple introgressions likely contribute to these variations. Nevertheless, the Siverskyi Donets center of water frog diversity remains rich in different forms of water frogs including polyploids and parental species. Moreover, year-to-year the portion of such forms remains stable. This makes it unique and distinct from Central European R-E systems initially described as “leaky all-male hybrid systems”, with triploids absent (Doležálková-Kaštánková et al., 2018; Uzzell et al., 1977).

### Genetic diversity and clonal lineages in the Siverskyi Donets center

In the R-E and R-E-Ep HPSs with the majority of diploid hybrids producing high portions of R- gametes, we would expect a high rate of clonality among R-genomes. In typical cases described in R-E HPSs from Central and Northern Europe, to successfully reproduce, most hybrids eliminate the R-genome premeiotically, transmit the L-genome to the gametes, and further cross with *P. ridibundus* (Graf & Polls-Pelaz, 1989; Plötner, 2005). However, in the Siverskyi Donets center, hybrids originate not only from crosses between *P. esculentus* and *P. ridibundus* but also from crosses involving *P. esculentus* alone, as hybrids here exhibit amphigameticity meaning they can produce a mixture of R- and L-gametes (Biriuk et al., 2016; Dedukh et al., 2017; Pustovalova et al., 2022a). Moreover, the rate of L-genome-carrying gametes is lower than that of R-genome-carrying ones (Biriuk et al., 2016; Dedukh et al., 2015; Fedorova & Shabanov, 2022; Pustovalova et al., 2024). We demonstrated that among all the studied R-genomes in R-E HPSs from Siverskyi Donets basin only five R-genomes formed two potential clonal lineages, while most of the L- genomes in hybrids clustered in at least 17 clonal lineages. Thus, the general clonality of R- genomes was significantly lower than that of L-genomes. The presence of many clonal lineages among L-genomes may indicate their multiple origins in the Siverskyi Donets center. In contrast, in the R-E system from Central Europe, where triploid hybrids are absent, L-genomes are presented as a single clonal lineage with extremely low genetic diversity, suggesting a single origin of L-genomes there (Doležálková-Kaštánková et al., 2018). In the Siverskyi Donets center, where both forms of triploids were found (Biriuk et al., 2016; Borkin et al., 2004, this study), the diversity of L-genomes is much higher. Since the recombination of doubled genomes was shown in triploids from Northern Europe (Christiansen & Reyer, 2009) we may also suggest potential recombination of L-genomes in LLR hybrids from Siverskyi Donets. Moreover, the pairing of L chromosomes was demonstrated in the meiosis of LLR males (Pustovalova et al., 2022a, 2024) and females (Dedukh et al., 2015) from studied localities.

We also observed a higher genetic diversity of R-genomes than of L-genomes in adult hybrids. The lower diversity of L-genomes in Siverskyi Donets basin results from the absence of *P. lessonae* as a main source of recombined genomes in this area. In the Dnipro basin, where *P. lessonae* is widespread, the diversities of L- and R-genomes were almost equal. The higher variability of R-genomes results from the presence of the sexual species *P. ridibundus*, which constantly adds recombined R-genomes to the gene pool. Potentially, LRR hybrids may also contribute to the pool of recombined R-gametes since the recombination of doubled genomes in triploids was shown previously for frogs from Northern Europe (Christiansen & Reyer, 2009). Even in HPSs with extremely low proportions of *P. ridibundus* and LRR hybrids, they can play a crucial role in maintaining R-genomes diversity. A similar situation was described in L-E HPSs from Central Europe, where *P. esculentus* producing R-gametes depend on very rare *P. lessonae* (Mikulíček et al., 2015).

Moreover, the uniqueness of Eastern Ukrainian water frogs was demonstrated previously by the distinct clusterization of R- and L-genomes in them compared with hybrids from different European population systems (Hoffmann et al., 2015). Additionally, Hoffmann et al. (2015) reported lower genetic diversity of L-genomes compared to R-genomes, which corresponds with our results for R-E HPSs in Siverskyi Donets basin.

### Selective mortality of progeny

Our analysis comparing the proportions of tadpole and adult forms revealed substantial numbers of *P. ridibundus* among tadpoles and froglets, but not among adults, in three HPSs (KOR, ISK, and DOB, Fig. 3). This higher prevalence of *P. ridibundus* among offspring aligns with the predominance of R-gametes produced by both male and female frogs in this region (Biriuk et al., 2016; Dedukh et al., 2015, 2017; Pustovalova et al., 2022a). The variance in *P. ridibundus* proportions is likely due to selective mortality of hybridolytic parental species (Günther & Plötner, 1988), as previously demonstrated in the *P. esculentus* complex within Northern and Central European population systems (Reyer et al., 2015; Vorburger, 2001). Moreover, we propose that differing wintering conditions, elevated levels of heterozygosity, and reduced parasite burdens may influence the selective survival of hybrids.

Deleterious mutations in clonally transmitted genomes in a homozygotes state can decrease the viability of the progeny (D. Shabanov et al., 2020; Vorburger, 2001). Alternatively, if the genetic diversity is high such deleterious mutations have a low chance to appear in the same homozygotic position and cause the death of the progeny. Here, we found high diversity of R-genomes in the tadpoles so the mortality of RR embryos is unlikely caused by genetic factors. Ecological conditions, such as continuous pond drying, could also influence the mortality of certain forms of frogs in the Iskiv pond HPS from studied localities (Drohvalenko et al., 2023). Similarly, the drying of ponds affect *P. ridibundus* froglets, which cannot pass wintering as they tend to hibernate underwater (Berger, 1982; Berger & Rybacki, 1994; Semlitsch & Reyer, 1992). Since adult frogs can produce gametes with the L-genome, we should have also expected to see *P. lessonae* tadpoles in the studied HPSs. However, no *P. lessonae* among both tadpoles and froglets was found. Their death could be explained by the presence of deleterious mutations in L-genomes, which are transmitted clonally (Reyer et al., 2015).

Previously a high frequency of morphological anomalies such as ectromelia, hemimelia, and taumelia was found among metamorphs from DOB (Fedorova et al., 2023). Such anomalies may reflect developmental disturbances usually found among hybridolytic progeny (e.g. Berger, 1968; Guex et al., 2002; Reyer et al., 2015). These anomalies impact frogs’ viability since among adult frogs their frequency was significantly lower (Fedorova et al., 2023; Kryvoltsevych et al., 2022). Unfortunately, in Fedorova et al. (2023) the species of froglets were not identified, so we cannot claim whether morphological anomalies play a role in the selective mortality of specific genotypes in the studied HPSs. Malformations can also signal changes in habitat conditions, any internal features of frogs, or be connected to infections, such as *Strigea robusta* (Svinin et al., 2023).

The selective mortality of some offspring is crucial for HPSs stability (Christiansen, 2009; D. Shabanov et al., 2020). Both water frogs HPSs and populations of sexual species include cycles of fertile individuals, gametes, zygotes, tadpoles, juveniles, immature individuals, and adults, with particular survival and mortality rates at each stage. In sexually reproducing species, such cycles lead to the transmission of the same genome throughout generations (Fu et al., 2019; Otto, 2008). However, in *P. esculentus*, hemiclonal genome transmission affects the genome composition of gametes, and thus shifts genotypes in the next generation of breeders (Som et al., 2000). Nevertheless, the long-term stability of some HPSs in the study area can be reached by the balance between constantly changing species composition and selective mortality of individuals due to environmental or genetic factors (D. Shabanov et al., 2020). If we assume that both species have equal viability, environmental factors could lead to the extinction of one form, disrupting the HPS or causing dominance by *P. ridibundus* (D. Shabanov et al., 2020; Som & Reyer, 2006, 2007). Therefore, to sustain the *P. esculentus*/*P. ridibundus* ratio in environmental disturbances, selection pressure should favour those genotypes whose proportion in the population system was affected (Som & Reyer, 2006, 2007). In addition to environmental factors, genetic factors also influence the development of individuals with specific genotypes. Such reduced viability of hybrid offspring is often linked to lethal mutations in clonal genomes (“Muller’s ratchet”) (Guex et al., 2002; Vorburger, 2001). However, it is still unclear why offspring viability is reduced, regardless of the clonal or recombined genomes inherited from the parents (Plötner, 2005). Thus, selective forces favour genomes that prevent the development of individuals with unfavourable genomic compositions, maintaining the *P. esculentus*/*P. ridibundus* ratios needed for HPS stability.

## Conclusion

In conclusion, our extensive investigation of water frog populations in the Siverskyi Donets River basin revealed diverse HPSs compositions, genetic divergence, and selective mortality patterns among tadpoles. We identified two main types of R-E systems: with diploid hybrids (R-E HPSs) and with diploid and triploid hybrids (R-E-Ep HPSs) extending beyond the mainstream of the Siverskyi Donets River to its tributaries. Eastern Ukrainian water frog population systems exhibit unique traits, including a rich diversity of triploid forms contributing to HPSs prosperity. Additionally, we observed higher genetic diversity in R-genomes compared to L-genomes, likely due to the continual input of recombined R-genomes from *P. ridibundus* and triploid LRR. Selective mortality of tadpoles within HPSs is evident, with a higher proportion of *P. ridibundus* among tadpoles compared to adult frogs. This phenomenon may be caused by deleterious mutations in hybridolytic parental species, ecological factors, or the development of morphological anomalies. The absence of *P. lessonae* tadpoles suggests early mortality due to mutations in L-genomes, while morphological anomalies among metamorphs may indicate developmental disturbances impacting frog viability.

Our study underscores the need for ongoing monitoring and research to comprehend the dynamics and intricacies of water frog population systems, critical for understanding factors influencing their genetic diversity, composition, and long-term survival.

## Author Contribution

AF, EP, MD, OB, DD, DS, PM, DH – Data Curation, Investigation, Methodology; OK, MK, MDK – Investigation, Methodology; AF, EP – Writing – Original Draft Preparation; LC – Methodology, Resources, DD, DS – Conceptualization, Resources, Supervision, Writing – Review & Editing; all the authors approved the final draft of the manuscript.

## Funding

EP was supported by the Academy of Sciences of the Czech Republic (RRFU-22-21) and EMBO (SLG-5411). AF was supported by the Academy of Sciences of the Czech Republic (RRFU-22- 20), EMBO (SLG-5409), and Slovak Academic Information Agency (1/0286/19). The work of DD, MDK, and AF was supported by the Czech Science Foundation grant (23-07028K). DD and MDK were also supported by the Czech Science Foundation grant RVO (67985904). M.D. was funded by a scholarship programme of the National Science Centre of Poland (UMO- 2022/01/4/NZ8/00020) for students and researchers from Ukraine who have taken refuge in Poland after the Russian invasion of Ukraine. The work of PM was supported by the Scientific Grant Agency of the Ministry of Education, Science, Research and Sport of the Slovak Republic and the Slovak Academy of Sciences VEGA (grant no. 1/0014/24).

## Conflicts of Interest

The authors declare no conflict of interest.

## Acknowledgments

Authors are very thankful to students, teachers, and friends who helped to perform the expeditions and catch the frogs, particularly during the students summer practice in 2012-2021. We also appreciate the incredible efforts of the Armed Forces of Ukraine whose bravery and sacrifice allow us to continue our scientific research.

## Notes

### Competing Interest Statement

The authors have declared no competing interest.

