## Supplementary Table S1 for "Hybridogenetic reproduction of *Pelophylax* water frogs from different R-E hemiclonal population systems from Eastern Ukraine: selective mortality, clonal and ploidy diversity"

| **Morphological trait** | **Pool frog — *Pelophylax lessonae* (Camerano, 1882)** | **Edible frog —*Pelophylax esculentus* (Linnaeus, 1758).** | **Marsh frog —*Pelophylax ridibundus* (Pallas, 1771)** |
| --- | --- | --- | --- |
| Size and form of metatarsal tubercle | Semicircular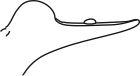 | Low, often semicircular, not sloped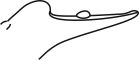 | Flat (low), sloped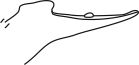 |
| Relative shin length | 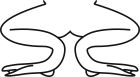 | 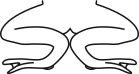 | 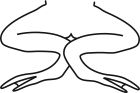 |
| Coloring of the dorsal surface of the body | Light green, grass green, or brown, ♂♂ during spawning can be lemon, with a few dark pigment spots | Light green or grass green, brown or bronze; numerous pigment spots are black, clearly defined | Olive-green with edging or brown with large brown (rarely greenish) bright, irregular spots |
| Dorsomedial band | Always present (*striata* form) | Always present (*striata* form) | Sometimes present (*striata* form), sometimes not (*maculata* form) |
| Coloring of the ventral surface of the body | White or slightly pigmented, rarely with gray spots or marble | White or gray-marble | From gray to blackish marble or spotted |
| Coloration of inguinal region underside of thighs | Intense yellow or warm yellow color inside and outside | Yellow spots often appear, especially during mating season; the pattern almost always has a warm yellow color | The back has whitish, grayish, or occasionally greenish spots, with cold sandy tones instead of warm yellow ones |
| Coloring of vocal sacs in ♂♂ | Always white, fully unpigmented | From white to dark gray (through all shades of gray) | From light gray to black |
| Smell | Weak | Strong | Pungent specific |
| Mating call of ♂♂ (after «chorus») | Chirping 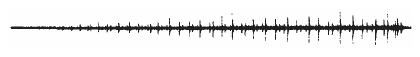 | Intermediate (similar to both parental species) 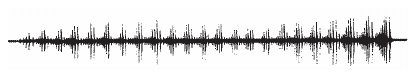 | Reverberating, laughter-like 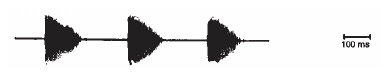 |
| Behavior in response to danger | Swim under water and emerge | Intermediate (similar to both parental species) | Hide at the bottom |
| ♂♂ behavior toward competitors during spawning and mating calls | Very aggressive  (attack, drive away, forced silencing) | Aggressive | Neutral |
| Wintering | On land | Both options, more often on land, often together with the co-existing parental species | Under the water |
| Preferred habitat | Small water bodies of the forest zone, outside the spawning period live on dry land | Different, except from extreme types common for the parental species | Large reservoirs of open landscapes |

**Fig. S1. Morphological traits for distinguishing between parental species (*Pelophylax lessonae*, *Pelophylax ridibundus*) and their hybrid (*Pelophylax esculentus*)** (Günther, 1975; Berger et al., 1978; Plötner 2005; Shabanov, 2015)
