## Supplementary Table S4 for "Hybridogenetic reproduction of *Pelophylax* water frogs from different R-E hemiclonal population systems from Eastern Ukraine: selective mortality, clonal and ploidy diversity"

**Table S3. Numbers of frogs possessing different multilocus genotypes (MLGs) in L- and R-genomes.** DOB – Lower Dobrytskyi pond, DVO – NNP "Dvorichanskyi", GRZ – Gruzotonovka, ISK – Iskiv pond, KOR – Koriakiv pond, LOZ – Lozovenky, LUH – Oleksiivskyi Luhopark, PIV – Pivdenne, PSL – Psel river, SLB – NNP "Slobozhanskyi", TYM – Tymchenky pond, UDAI – Udai river, UDY – Udy river, VOD – Vodiane, ZAM – Zamulivka village, ZHK – Zhykhorets river, ZHR – Zhuravlivskyi Hydropark.

| **MLG** | **N of frogs** | **River basin** | **Localities** |
| --- | --- | --- | --- |
| *L-genomes* |  |  |  |
| A | 3 | Siverskyi Donets | ZHR, KOR |
| B | 18 | Siverskyi Donets | ZHK, DOB, KOR, ZHR, DVO |
| C | 2 | Siverskyi Donets | KOR |
| D | 6 | Siverskyi Donets | VOD |
| E | 9 | Siverskyi Donets | ZHK, ZHR, DOB, KOR |
| F | 2 | Siverskyi Donets | ZHR, KOR |
| G | 19 | Siverskyi Donets | ZHK, PIV, DOB, DVO, VOD, KOR, TYM |
| H | 5 | Siverskyi Donets | ZHK, DOB, KOR |
| I | 7 | Siverskyi Donets | DOB, TYM, KOR |
| G | 31 | Siverskyi Donets | VOD, PIV, GRZ, UDY, LOZ, TYM, KOR |
| K | 2 | Siverskyi Donets | ZHK, DOB |
| L | 5 | Siverskyi Donets | PIV, UDY |
| M | 2 | Siverskyi Donets | ZHK |
| N | 6 | Siverskyi Donets | ZHK, GRZ, DOB, UDY, TYM |
| O | 5 | Siverskyi Donets | LUH, UDY, TYM |
| Unique | 37 | Siverskyi Donets  Dnipro | KOR, LOZ, PSL, SLB, UDAI, VOD, VRG, ZHR, ZHK |
| *R-genomes* |  |  |  |
| A | 2 | Siverskyi Donets | DOB |
| B | 2 | Siverskyi Donets | VOD, KOR |
| C | 2 | Siverskyi Donets | VOD |
| D | 2 | Siverskyi Donets | ZHK, TYM |
| E | 4 | Dnipro | SLB |
| Unique | 173 | Siverskyi Donets  Dnipro | DOB, DVO, GRZ, KOR, LOZ, LUH, PIV, PSL, SLB, TYM, UDY, VOD, VRG, ZHR, ZHK |
